# Spatial transcriptomics reveals novel genes during the remodelling of the embryonic human arterial valves

**DOI:** 10.1101/2023.05.09.539950

**Authors:** Rachel Queen, Moira Crosier, Lorraine Eley, Janet Kerwin, Jasmin E. Turner, Jianshi Yu, Tamil Dhanaseelan, Lynne Overman, Hannah Soetjoadi, Richard Baldock, Jonathon Coxhead, Veronika Boczonadi, Alex Laude, Simon J. Cockell, Maureen A. Kane, Steven Lisgo, Deborah J. Henderson

## Abstract

Abnormalities of the arterial valves, including bicuspid aortic valve (BAV) are amongst the most common congenital defects and are a significant cause of morbidity as well as predisposition to disease in later life. Despite this, and compounded by their small size and relative inaccessibility, there is still much to understand about how the arterial valves form and remodel during embryogenesis, both at the morphological and genetic level. Here we set out to address this in human embryos, using Spatial Transcriptomics (ST). We show that ST can be used to investigate the transcriptome of the developing arterial valves, circumventing the problems of accurately dissecting out these tiny structures from the developing embryo. We show that the transcriptome of CS16 and CS19 arterial valves overlap considerably, despite being several days apart in terms of human gestation, and that expression data confirm that the great majority of the most differentially expressed genes are valve-specific. Moreover, we show that the transcriptome of the human arterial valves overlaps with that of mouse atrioventricular valves from a range of gestations, validating our dataset but also highlighting novel genes, including four that are not found in the mouse genome and have not previously been linked to valve development. Importantly, our data suggests that valve transcriptomes are under-represented when using commonly used databases to filter for genes important in cardiac development; this means that causative variants in valve-related genes may be excluded during filtering for genomic data analyses for, for example, BAV. Finally, we highlight “novel” pathways that likely play important roles in arterial valve development, showing that mouse knockouts of RBP1 have arterial valve defects.

Thus, this study has confirmed the utility of ST for studies of the developing heart valves and broadens our knowledge of the genes and signalling pathways important in human valve development.

**Non-Technical Summary:** Congenital heart defects, particularly those affecting the valves and septa of the heart, are very common. Despite this, few gene variants have been confirmed as disease-causing in human congenital heart (including valve) disease patients. Here we utilise spatial transcriptomics technology, which allows the identification of genes expressed in tissue slices, on embryonic human heart valves and identify a gene dataset that is human arterial valve-specific. We confirm the localisation of key novel genes to the arterial valves and highlight the relevance of the dataset by showing that mice mutant for RBP1, a novel gene identified as being highly differentially expressed in our valve dataset, have previously unidentified arterial valve defects. Using commonly used bioinformatic databases we show that filtering patient genomic data using these terms would likely exclude valve genes and thus may not identify the causative genes. Thus, we confirm that spatial transcriptomics technology can be used to study gene expression in tiny structures such as the developing heart valves and provide a new human embryonic valve dataset that can be used in future genomic studies of patients with congenital valve defects.

## Introduction

Abnormalities in the cardiac valves are amongst the most common congenital birth defects and cause significant morbidity throughout the lifespan. Over 1 in 100 of the population have a bicuspid aortic valve (BAV). Most are asymptomatic until adulthood when progressive fusion of the leaflet commissures causes clinically apparent aortic stenosis (AS). In addition, accelerated calcification and aneurysmal dilatation of the proximal aorta occurs in many BAV patients (Siu and Silversides, 2010). Fetal aortic stenosis is usually progressive and, at birth, may present in isolation as critical AS, or form part of complex lesions, for example, hypoplastic left heart syndrome (Brenner, 1989). Although in isolation BAV is often asymptomatic, it is frequently associated with more serious lesions and predisposes to severe cardiovascular disease in later life. For this reason, it has received considerable attention in recent years.

Although a number of large-scale genomic studies have been carried out to look for disease genes in patients with congenital heart defects (CHD) (Zaidi et al, 2013; Homsy et al, 2015; Glessner et al, 2014) and more specifically BAV (Mohammed et al, 2006; McKellar et al, 2007; Wooten et al, 2010; Martin et al, 2014; Bonachea et al, 2014; Dargis et al, 2016; Giusti et al, 2017; Yang et al, 2017 Gharibeh et al, 2018), few genes have been identified, and even fewer gene variants have so far been verified as specifically causing BAV. In those cases where genes have been suggested to be causative for BAV, the whittling down of huge numbers of patient variants to a handful of gene candidates has been carried out by reference to datasets of genes known to be important in cardiac development – heavily reliant on developmental genetic studies in animal models (for example Mohammed et al, 2006; McKellar et al, 2007; Yang et al, 2017; Gharribeh et al, 2018). Thus, identification of genes as being expressed in the developing mouse heart, followed by functional analysis of these genes in animal models, remains the cornerstone of detection of human CHD and BAV causative genes. Arguably, however, the absence of valve-specific gene expression datasets is hampering this approach and explains, at least in part, the difficulties experienced in confirming gene variants as causative for BAV.

Although aortic valve disease is an area of major clinical importance, most of what we know is derived from analysis of the atrioventricular valves. Only recently have studies focussed specifically on the differential development of the arterial valves (Eley et al, 2018; Mifflin et al, 2018, Peterson et al, 2018; Milos et al, 2018; Henderson et al, 2020). Consequently, while our knowledge of the formation of the cushions, the precursors for the mature valve leaflets is relatively well established, comparatively little is known about how these bulky structures remodel into sculpted valve leaflets. Studies in zebrafish have suggested that haemodynamic forces resulting from blood flow through the heart are critical for this remodelling process (Kalogirou et al, 2014; Steed et al, 2015; Goddard, et al, 2017), but how they integrate with intrinsic, genetically regulated, signalling cascades remains unclear. Notably, there remains a specific lack in our current understanding about the development of the human arterial valve leaflets (Kramer 1942; Maron and Hutchins, 1974; Henderson et al, 2022), although it is generally assumed that the morphogenetic processes will be similar to those implicated from the study of animal models. Although this is likely correct, there is some evidence for different patterns of aortic valve anomalies in mouse and human, indicating that the reliance on particular processes or susceptibility to pathology may vary between the two species (discussed in Henderson et al, 2020).

Single cell transcriptomics (sc-RNASeq) has become commonplace over recent years, with increasing evidence that this approach can bring new insights to normal physiology and disease pathology (Jardine et al, 2021). Here we utilised a spatial transcriptomic approach (Stahl et al, 2016; DeLaughter et al, 2016; Vickovic et al, 2019) to derive data about gene expression in the arterial valves of the human embryo, during the sculpting phases of their development. We have been able to validate these gene sets by comparison with sc-RNAseq and in some cases spatial transcriptomic datasets from other studies of fetal and postnatal valves, as well as by confirmation through RNA in situ expression experiments in mouse and human embryos. We developed bioinformatic approaches for transcriptomic analysis, which were used to identify a number of genes that have not been previously linked to valve development and are strongly expressed in the remodelling valve leaflets. Examination of the mouse model for one of these, RBP1, shows previously undescribed defects in the great arteries and valve anomalies in mutant animals, highlighting that these genes are good candidate genes for causing BAV. Through the use of specialised spatial analysis methodologies, we show ST datasets have the ability to identify previously unknown developmentally important genes. We have shown that ST can be used as an powerful technique for novel gene discovery in the developing human embryo.

## Results

### Optimisation of spatial transcriptomics for analysis of developing human heart valves

There have been few transcriptomic analyses of developing heart valves, particularly in the human embryo. To circumvent the problem of dissecting out these tiny structures in this scarce material, we utilised ST technology to bioinformatically distinguish the forming valves from the surrounding myocardial tissue.

Dissected and cryo-embedded chromosomally normal CS16 (approximately 37 post conception days) and CS19 (approximately 48 post conception days) hearts, both from female embryos, were chosen for analysis, as these map to the period of development when the arterial valves are beginning to remodel from bulky cushions to more sculpted valve leaflets - a time frame where relatively little is known (Figure 1A). Previous studies from our laboratory have shown that the three leaflets of the developing arterial valves have distinct origins (Phillips et al, 2013; Eley et al, 2018; Henderson et at 2020) and for this reason we elected to section the hearts in a cross-sectional plane through the aortic and pulmonary valve leaflets so that each of the three leaflets would be included in the analysis. H&E staining confirmed that the three valve leaflets could be readily distinguished in these hearts (Figure 1B).

**Figure 1.**
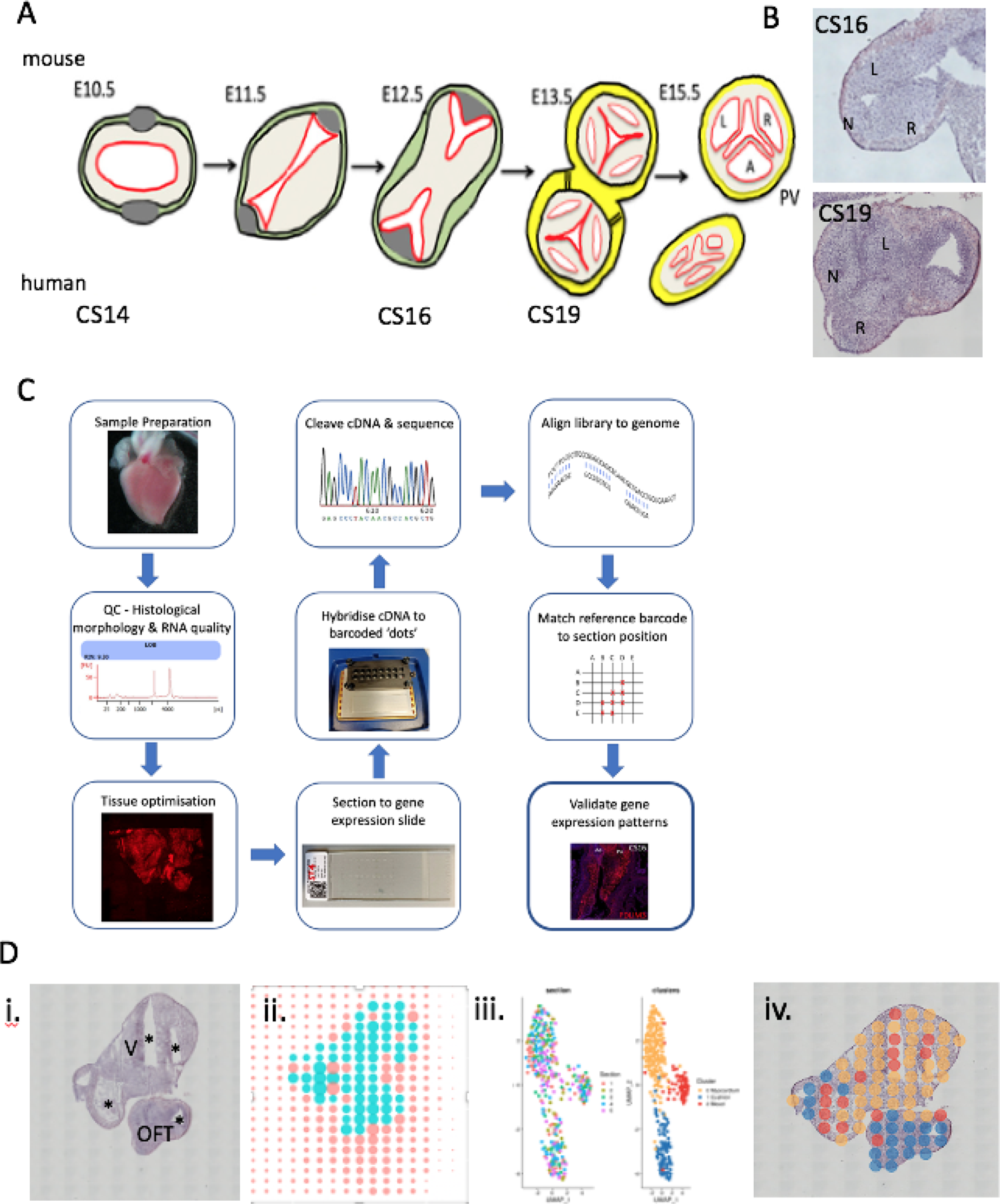
Overview of ST process. **A)** Cartoon showing stages of arterial valve development in mouse with human equivalents. **B)** H&E -stained sections of CS16 and CS19 human arterial valves used for analysis. **C)** workflow of ST including quality control steps. **D)** Example of output of ST for CS19 embryo: i) H&E section; ii) position of spots on hybridisation slide with turquoise spots covering tissue section and orange spots covering blank slide; iii) the first graph shows that data from each of the 5 slides (each with a different colour) from the CS19 embryo slide overlaps following clustering, whilst the second graph shows the three clusters (yellow, red, blue) obtained; iv) three clusters mapped back onto H&E sections. L = left leaflet; N = non-coronary leaflet; OFT = outflow tract; R = right leaflet; V = ventricle; * lumen of chambers and/or outflow tract.

An overview of the adopted experimental design is outlined in Figure 1C and follows a previously described method (Asp et al, 2017) with the addition of quality control checks prior to the beginning of the workflow and following bioinformatics analysis, as well as supplementary data analysis which involved positioning of the transcripts to digitally annotated anatomical regions. On average there were more than 6400 counts and 2200 genes found in each spot at CS16, and 17000 counts and 3700 genes per spot at CS19. Cluster analysis of the technical replicates from each time point individually resulted in three clusters when a resolution of 0.8 was used (Figure 1D). Clustering of spots was not section-specific and spots from all sections at a particular stage were found throughout the dataset (Figure 1D). This indicates that the ST is reproducible and shows that clusters identified are not driven by technical factors in the data.

### The transcriptomes of CS16 and CS19 aortic valves are comparable

To further confirm that the transcriptomic data obtained were biologically reproducible, we compared the datasets obtained at CS16 with that obtained at CS19 and visualised them on a UMAP. These analyses showed that, similar to Asp et al (2019), the two datasets overlapped, with no obvious separation between them (Figure 2A). Despite the difference in developmental maturity between the two samples, their transcriptomes were highly comparable. This allowed the two samples to act as biological controls for one another and for this reason, all further analyses were carried out using a combined CS16/19 dataset.

**Figure 2.**
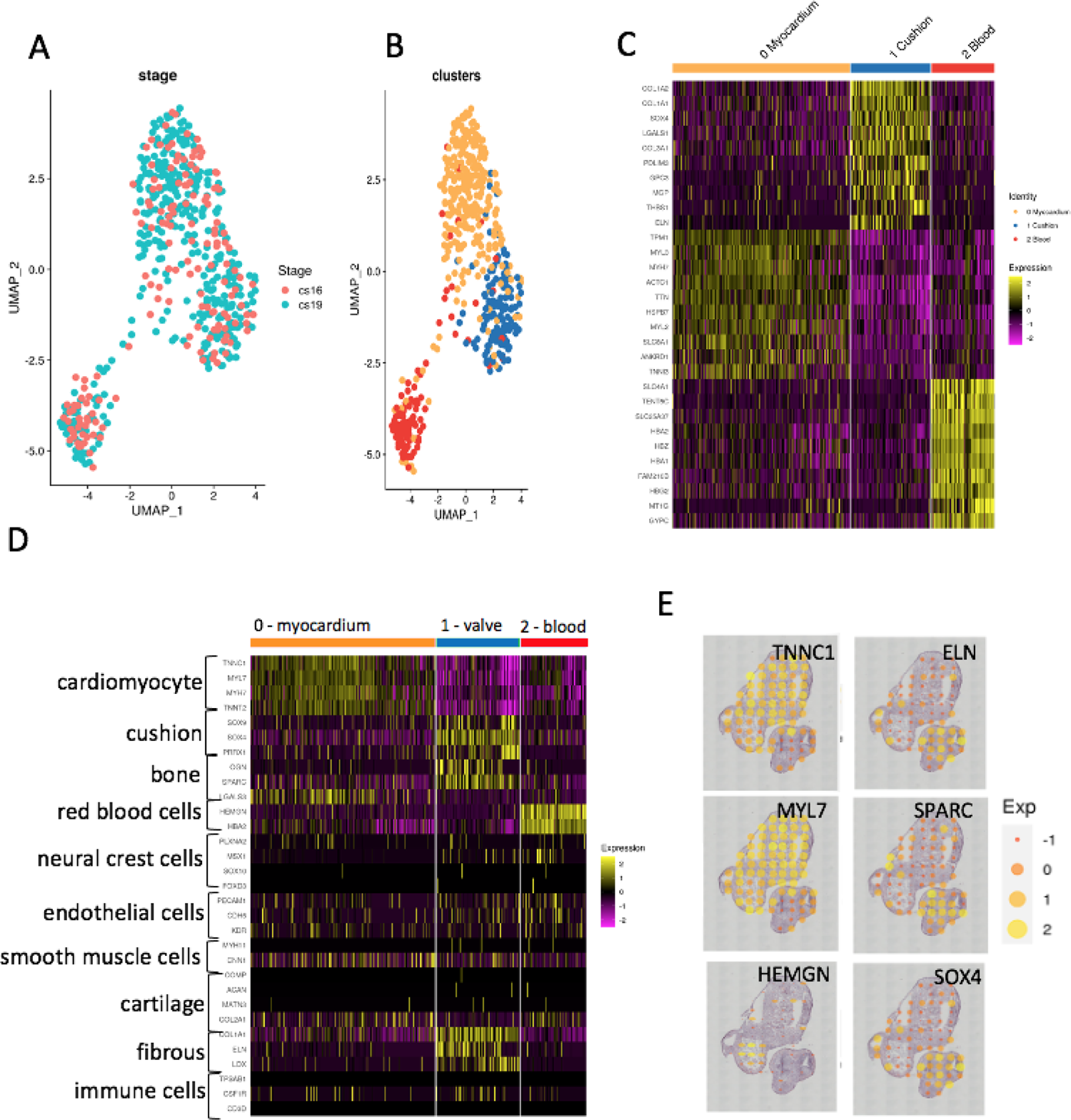
Cluster analysis. **A)** UMAP projection showing overlap of data between sections from the CS16 (orange dots) and CS19 (turquoise dots) embryos. **B)** UMAP projection showing clustering of combined CS16 and CS19 data, again showing three clusters (red, blue and orange dots). **C)** Heat map showing top 10 genes in each cluster, highlighting the gene expression differences between the three clusters. At this stage, based on the expressed genes, putative tissue types can be proposed as myocardium (orange), cushion (blue) and blood (red). **D)** Perfect marker analysis using recognised markers for specific cardiac cell types confirms the orange cluster as myocardium, the red cluster as blood cells, and shows that the putative cushion cluster has characteristics of cushions, bone and fibrous tissue, and thus should be better named as “valve” as this implies the entire structure, not just the developing leaflets. **E)** Mapping “perfect markers” back to the H&E sections shows that they localise to the expected areas (compare to Figure 1D).

The differentially expressed genes at CS16/19 fell into three clusters (Figure 2B), with heatmaps of the most highly differentially expressed genes confirming clear separation (Figure 2C). “Perfect Marker” analysis, using marker genes identified in previous studies as being characteristic of specific differentiated cell types in the heart (for example deLaughter et al, 2016; deSoyza et al, 2019; Asp et al, 2019), confirmed two of the three clusters as cardiomyocytes (cluster 0) and blood cells (cluster 2). The relevant “Perfect Marker” genes mapped back to the expected regions of the histological sections at both time points (Figure 2E and Supplementary Figure 2). The third putative cluster (cluster 1) was more complex than the others, with genes characteristic of endocardial cushion tissue, bone and fibrous tissue. As endocardial cushions (the precursors of the valve leaflets) have been described previously to share many markers with developing bone (Chakraborty et al, 2008; see below) and the walls surrounding the maturing cushions/valves become progressively fibrous (Richardson et al, 2018) this cluster likely corresponds to the developing valve complex. Moreover, the relevant marker genes mapped back to the arterial valve region on the tissue sections (Figure 2E and Supplementary Figure 2). Interestingly, neural crest cell marker genes were not abundant in the tissue, suggesting that these progenitor cells had differentiated by CS16, nor were genes typically expressed in smooth muscle cells or cartilage. Typical endothelial cell genes were evenly distributed between the three clusters, reflecting the abundance of this cell type in all tissues (Figure 2D and Supplementary Figure 2).

### Novel aortic valve-specific genes in human aortic valves

The top 30 differentially expressed genes in the combined CS16/CS19 dataset, for each of the three clusters, are shown in Figure 3A (For full gene lists see Supplementary Tables 1-3; GEO – submission in process). Genes that have been previously identified as being strongly expressed in cardiomyocytes, cushions/valves or blood are colour coded as for the cluster diagrams, whereas those previously not known to be expressed in the relevant tissues (although they may be known to be expressed in the heart in general) are denoted in black. Analysis of the genes in these clusters supports the idea that cluster 0 represents cardiomyocytes, with several members of the myosin light and heavy chain, troponin and tropomyosin families most highly expressed; 29 genes in the top 30 had previously been shown to be highly expressed in cardiomyocytes. In the case of the putative “cushion/valve” cluster (cluster 1), the most highly expressed genes have previously been associated with the developing arterial wall that surrounds the developing valve complex (including COL1A1, COLIA2, COL3A1 and ELN), as well as genes associated with the developing cushions such as SOX4. This supports the findings of the “perfect marker” analysis, which showed the presence of fibrous markers known to be abundant within the arterial wall associated with this cluster. It would therefore seem more appropriate to characterise this as a “valve” cluster as this more general term would include the cushion-derived leaflets as well as the supporting arterial structures. Cluster 2 was confirmed as blood cells within the lumen of the heart and includes an abundance of genes known to be expressed in erythrocytes including multiple haemoglobin isoforms (Figure 3A).

**Figure 3.**
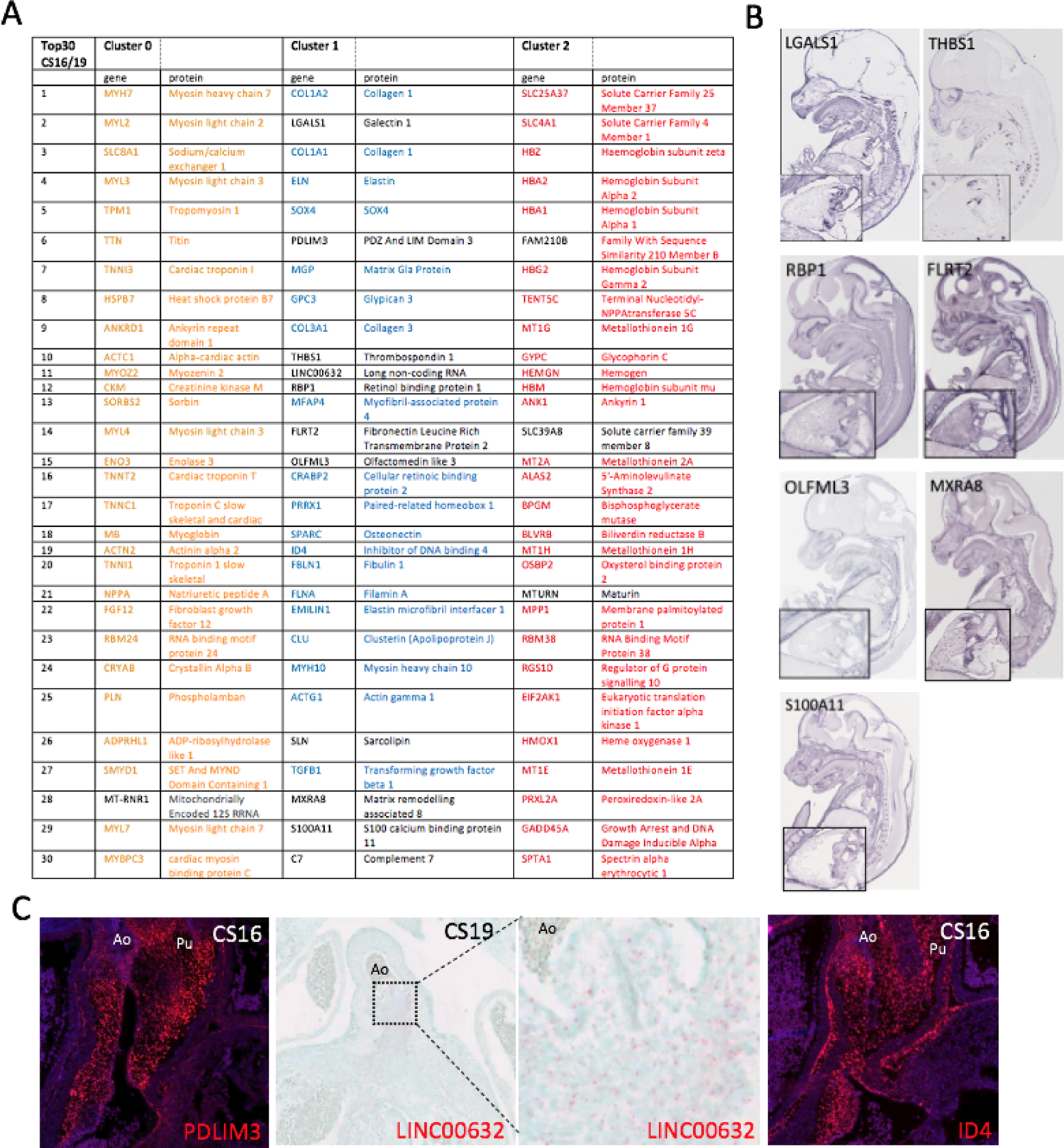
Identification of cluster genes. **A)** Top 30 differentially expressed genes in each of the 3 clusters. In each case, the coloured genes are already documented to be expressed in the proposed cluster tissue, whereas genes in black font were not identified as being expressed in the relevant tissue by literature search. **B)** Expression of novel Cluster 1 (valve) genes in the developing E14.5 mouse heart (GenePaint images). **C)** PDLIM3, LINC00632 and ID4 transcripts (red dots in all cases) are all found in the CS16/CS19 human arterial valves. Ao = aortic valve; Pu = pulmonary valve.

We focussed on the valve cluster which was the main interest of our study. The majority of the top 30 differentially expressed genes had been previously associated with valve tissues, with analysis in the GenePaint mouse gene expression database (gp3.mpg.de) revealing remarkable valve-specificity (Figure 3B and Supplementary Figure 3). However, for 11/30 of the genes: LGALS1, PDLIM3, THBS1, LINC00632, RBP1, FLRT2, OLFML3, SLN, MXRA8, S100A11 and C7 there were, to the best of our knowledge, no published papers describing functions within the developing outflow tract, including the arterial valves. Of these “novel” valve genes, GenePaint showed that LGALS1, THBS1, RBP1, FLRT2, OLFML3, MXRA8 and S100A11 were all expressed in the arterial valve region in the E14.5 mouse, with no information available for PDLIM3 and LINC00632 (Figure 3B). SLN and C7 were represented in GenePaint but there was no clear localisation to the aortic valve (Supplementary Figure 3). We therefore carried out RNAScope and BaseScope gene expression analysis to investigate the expression pattern of PDLIM3 and LINC00632 respectively, as well as ID4 which is known to be expressed in the heart but lacked relevant expression data in GenePaint. All three genes localised to the aortic root at CS16 and CS19 (Figure 3C and Supplementary Figure 3B). Thus, the ST method we utilised has proven useful for identifying genes previously unknown to be involved in arterial valve development.

### The developing human aortic valve transcriptome is similar to mouse

We compared our data with other published datasets; it should be noted that the available data varied between studies and this influenced the depth of the comparisons that could be made. Whilst there were no datasets that were directly parallel either in human or mouse (where E12.5-E13.5 best matches the CS16/19 samples we used), a comparison with the top 250 differentially expressed genes between E11 mouse AV/OFT compared to ventricle (DeLaughter 2013) and our top 250 revealed a 20% overlap (Supplementary Table 4). Better overlap was obtained with a spatial transcriptomics dataset from 4.5-9 weeks post conception human hearts (Asp et al, 2019) where the figure was closer to 30% for a comparison between the AV cushions/valves cluster for the top 250 genes. Comparing the top 20 genes, five genes (25%; LGALS1, PDLIM3, MFAP4, PRRX1 and ID4) were common to both datasets Figure 4A). Comparison with the outflow tract/large vessel cluster from the Asp dataset revealed 70/250 overlapping genes and two (ELN, CRABP2) were shared between the top 20 from both datasets (Supplementary Table 4). We next compared our data with single cell RNAseq data from aortic/mitral valves from postnatal day (P) 7 and P30 mouse (Hulin et al, 2019). Although there was little overlap between our dataset and valve endothelial cells, valve immune cells or melanocytes, there was remarkable overlap with the most highly expressed genes in valve interstitial cells (VIC) (Figure 4A), which made up more than 75% of the cells in their dataset. COL1A1, COL1A2, LGALS1, MGP, COL3A1, FBLN1 and SPARC were in the top 20 differentially expressed genes in both datasets. Of Interest, one gene, LGALS1 was found in the top 20 differentially expressed genes in ours, the Asp (AV mesenchyme and valves) and the Hulin datasets. Expression analysis in mouse and human using an LGALS1-specific antibody showed that it was strongly expressed in the developing cushions in both species at E11.5/CS16, although the expression was more restricted to the forming valve region in the human embryo than in the mouse (Figure 4B). Finally, we compared our dataset to transcriptomic and protein profiling comparing non-diseased, fibrous and calcific aortic valves from human patients (Schlotter et al, 2018). There was almost no overlap with the transcriptomic datasets, however, there was overlap between the Schlotter proteomic data and our transcriptomic data, with 5/40 overlapping molecules in the non-diseased samples (COL1A2, COL1A1, FLNA, EMILIN1 and MYH10) and 5/40 in the fibrotic valve samples ((OGN, LGALS1, MGP, FBLN1, S100A10) – compared to our top 40 differentially expressed genes (Supplementary Table 4). There was no overlap between our dataset and the differentially expressed proteins in the calcific valve samples. Thus, our dataset validates previous microarray/RNASeq experiments carried out with mouse and human valve tissue, adding a significant number of “novel” genes – including LGALS1 - not previously identified as being important in developing arterial valve tissue.

**Figure 4.**
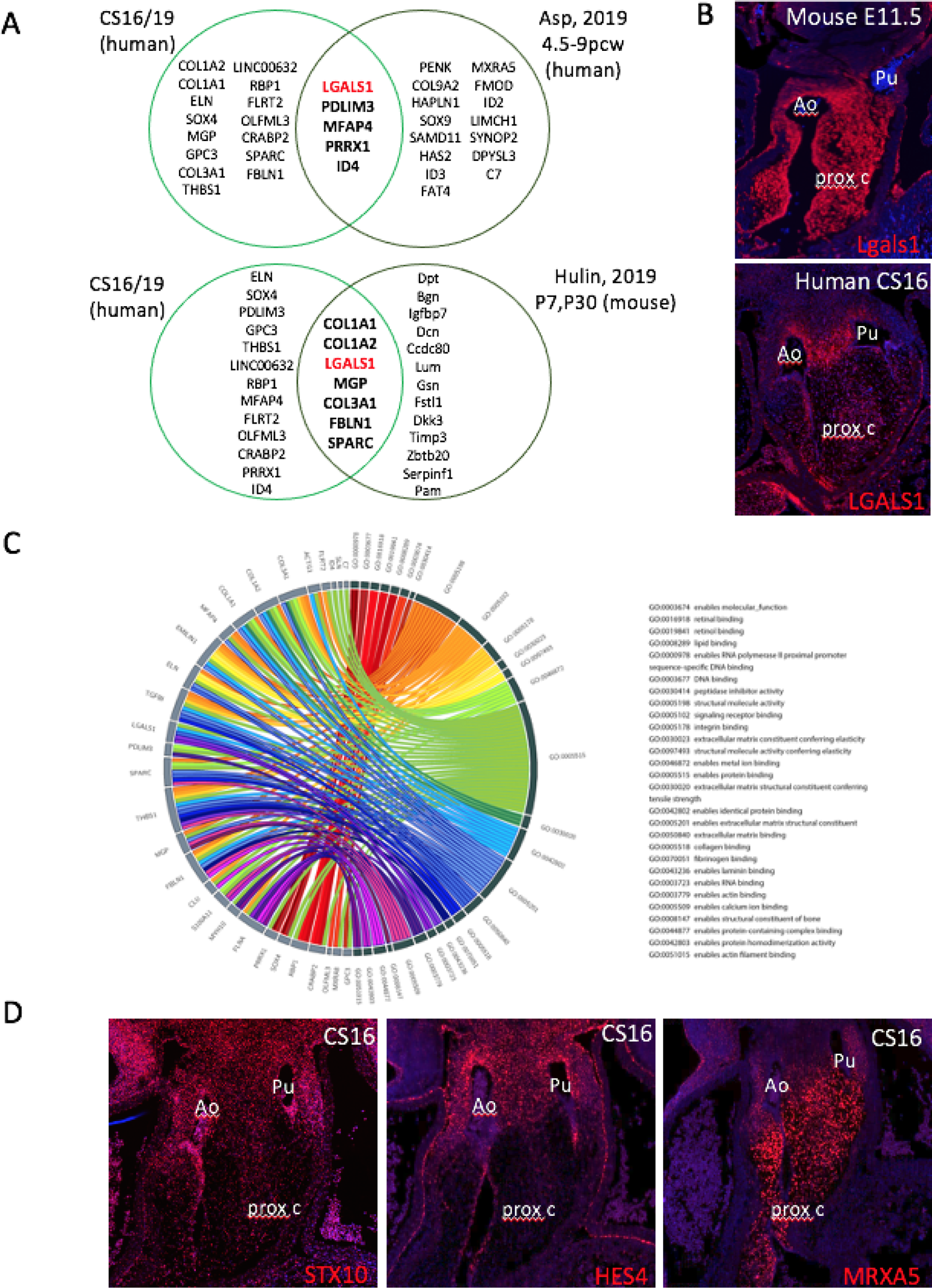
Comparison of “valve” genes to published ST/single cell RNASeq datasets. **A)** Comparison of the top 20 genes in our valve cluster with the human embryonic “atrioventricular mesenchyme and valve” ST cluster from Asp et al (2019) reveals 5 shared genes. Similarly, comparison of our top 30 valve cluster genes with scRNASeq “VIC” cluster data from P7 and P30 mouse atrioventricular and arterial valves (Hu,in et al, 2019) reveals 7 genes in common. Notably, LGALS1, which has not previously been reported to be specific to the arterial valves, is found in all 3 datasets. **B)** Immunohistochemistry for LGALS1 shows that it is expressed in E11.5 mouse and CS16 outflow cushion mesenchyme, although whereas it is widespread throughout the proximal cushions and the forming aortic (Ao) and pulmonary (Pu) valves in the mouse, it is more restricted to the valve forming region in human embryos. **C)** Circos plot showing the top30 genes in our valve cluster mapped to GO terms. **D)** STX10, HES4 and MRXA5, none of which are found in the mouse genome, are expressed (red dots) in the developing outflow tract, localising to the walls of the forming valve complex (STX10 and HES4) or to the cushion/leaflet mesenchyme (MXRA5). Ao = aortic valve; prox c = proximal cushions; Pu = pulmonary valve.

We analysed the top 30 genes identified in our combined CS16/19 valve dataset in GORILLA (http://cbl-gorilla.cs.technion.ac.il/), which allowed us to determine the molecular function of the genes. The most common functions were “enables protein binding”, “structural molecule activity” and “enables extracellular matrix structural constituent” (Figure 4C). The three transcriptional regulators in our top 30 most highly expressed genes in the combined CS16/19 dataset were SOX4, PRRX1 and ID4 (Figure 3A and 4C). We also analysed the top 250 differentially expressed genes identified in our combined CS16/19 valve dataset in PANTHER (http://www.pantherdb.org), which allowed us to determine the protein class for the genes. The most abundant categories were cytoskeletal proteins, extracellular matrix molecules, gene specific transcriptional regulators and metabolite interconversion enzymes (Supplementary Figure 4). We also looked at the biological processes implicated by our top 250 valves genes, again using PANTHER. This suggested that genes involved in “cellular processes”, “biological regulation” and “metabolic processes” were most abundant (Figure 4C and Supplementary Figure 4).

Of particular interest, four genes in the top 250 (STX10, HES4 and MXRA5, as well as LINC00632; described earlier) were not found in the mouse genome. RNAscope analysis revealed that like LINC00632 (Figure 3C), STX10, HES4 and MXRA5 were all expressed in the developing valve complex (Figure 4D). However, whereas STX10 and HES4 appeared to be largely restricted to the walls of the forming arterial roots, MXRA5 was found in the endocardial cushions that will give rise to the valve leaflets. Thus, we have identified genes highly expressed in the developing human valve complex that would not be picked up in transcriptomic studies in mouse embryos.

### Pathway analysis suggests valves genes under-representation of valve transcriptomes in IPA

We carried out Ingenuity pathway analysis (IPA) on the valve cluster data to further link our data into pathways and processes. We first looked at the predicted upstream regulators associated with our dataset (Figure 5A and Supplementary Figure 5). This suggested CXCL12 and TGFBR1, both of which are implicated in cushion/valve development (Ridge et al, 2021; Wang et al, 2006) as the most important regulators. Several Epithelial Mesenchymal Transition (EMT) regulators including SNAI2, CTTNB1, SNA1 and TWIST were also predicted to be important (Figure 5A). We linked the top 30 regulators to biological functions and found that tumorigenesis, cell movement and cell proliferation were the top processes, followed by cytoskeleton and cardiovascular disease (Figure 5B). It is perhaps unsurprising that these were the most prominent functional terms as both tumorigenesis and cushion formation share the common processes of EMT, cell migration and proliferation.

**Figure 5.**
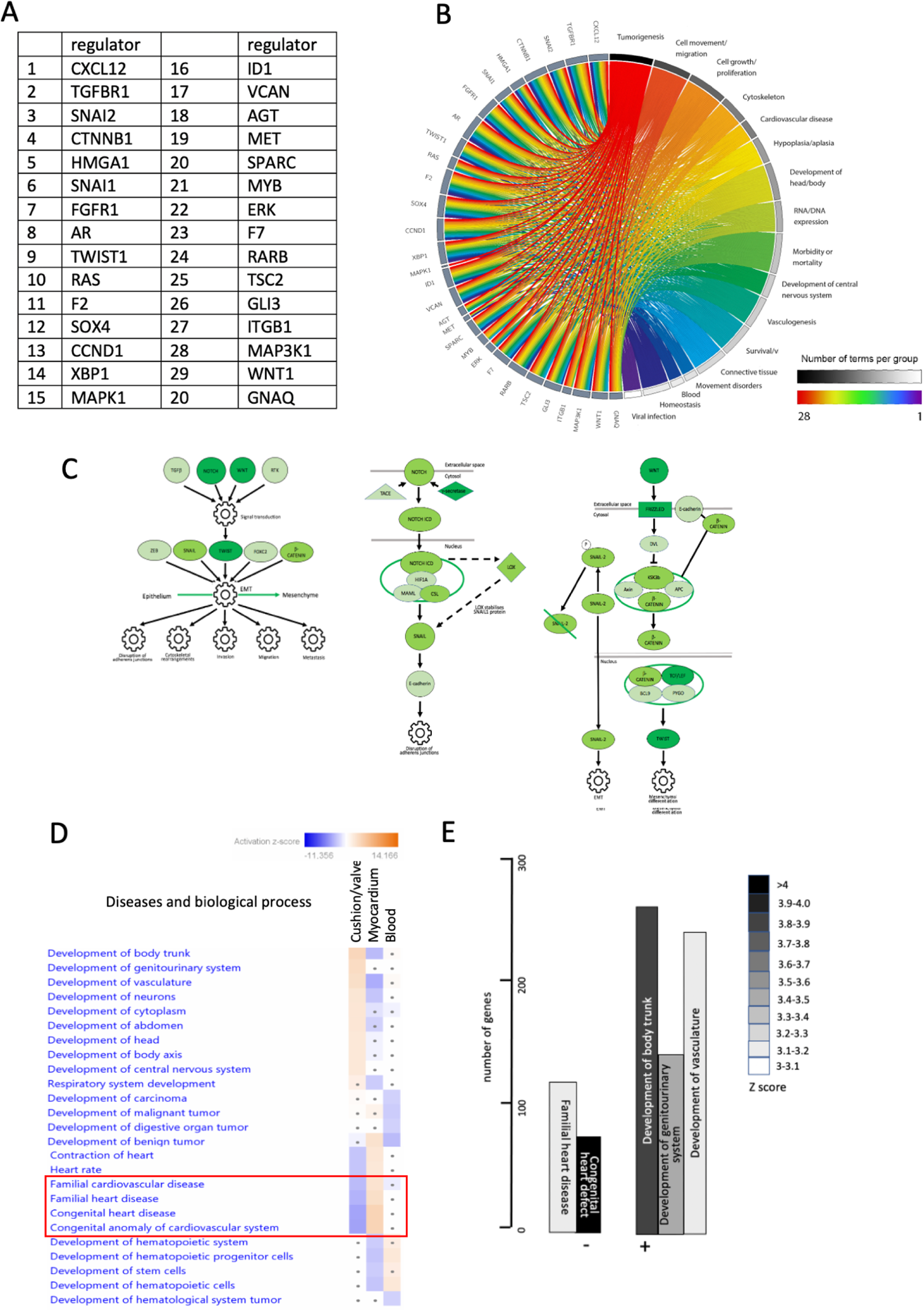
Functional associations of valve cluster genes generated by filtered IPA analysis. **A)** Top 20 regulators of genes in the valve cluster. **B)** Circos plot linking valve regulators to cellular processes. The strongest associations relate to cell movement and growth/proliferation but cardiovascular disease is also a string association for this group of valve regulatory genes. **C)** Following filtering for the terms “cardiac” and “development” regulation of EMT was the only pathway significantly upregulated in the valve dataset compared to the myocardial and blood clusters. Notch and Wnt were the likely upstream regulators. Pathways highlight the genes in the Notch and Wnt pathways that are positively associated regulators of EMT for the valve cluster (dark green genes are the most differentially upregulated, with mid green and light green progressively less so). **D,E)** IPA pathway analysis shows that for disease and biological processes, developmental terms such as development of trunk, genitourinary system, vasculature and neurons were strongly associated with the valve cluster. Negative associations were also found, the most striking being those to familial heart/cardiovascular disease, and congenital heart/cardiovascular disease/anomaly. These terms were positively associated with the myocardial cluster dataset (red box).

We then filtered our gene set for pathways associated with cardiac and development (using the terms card* and dev*), from the combined datasets from both time points (Figure 5 and Supplementary Figure 5C). Only the Regulation of the EMT in Development Pathway was significantly upregulated (more than three-fold compared to the myocardial and blood clusters) in the valve cluster. The data suggested NOTCH and WNT may be regulating the process of EMT, via SNAI2, CTTNB1, SNA1 and TWIST (Figure 5C and Supplementary Figure 5B). Developmental terms (trunk, genitourinary and vasculature) were all significantly positively associated with the valve cluster using the cardiac development filtering protocol. However, terms associated with “Congenital Anomaly of Cardiovascular System/Congenital Heart Disease” and “Familial Cardiovascular/Heart Disease” were significantly negatively associated (Figure 5 D,E), although they were positively associated with the myocardial cluster.

### Novel pathways associated with arterial valve development

In view of the single significant positively associated pathway (EMT), and the surprising negative association between the valve cluster genes and terms related to congenital heart disease, we carried out a second analysis, without filtering for cardiac and development, but otherwise using the same significance cut off of +/- two-fold compared to the myocardial and blood clusters. In this case, The HOTAIR Regulatory Pathway and EIF2 Signalling were the top two canonical pathways associated with the valve cluster (Figure 6A and Supplementary Figure 6 A,B). RNAScope analysis confirmed that HOTAIR and EIF2A were expressed in the endocardium of the developing human aortic valve (Figure 6B). GNRH Signalling, Ephrin Receptor Signalling and Regulation of Epithelial to Mesenchymal Transition were also significantly upregulated in arterial valve tissue. In the absence of filtering, the positively associated biological process and disease terms were involved in cell movement/migration and viability/survival/proliferation (Figure 6 C,D and Supplementary Figure 6C). This fits well with the dynamic cell movements and rapid proliferation that is taking place in developing valve tissues at these developmental stages (reviewed in Henderson et al, 2020). Negatively-associated terms were linked to growth failure or cell death, and similar to the filtered data, with familial heart disease, congenital heart defect and valvulopathy, although again, these were highly positively correlated in the myocardial cluster (Figure 6 C, D and Supplementary Figure 6C). Notably, valvulopathy was also negatively correlated with our cushion/valve cluster, supporting the lack of correlation with valve calcification datasets (Schlotter et al, 2018) as discussed earlier.

**Figure 6.**
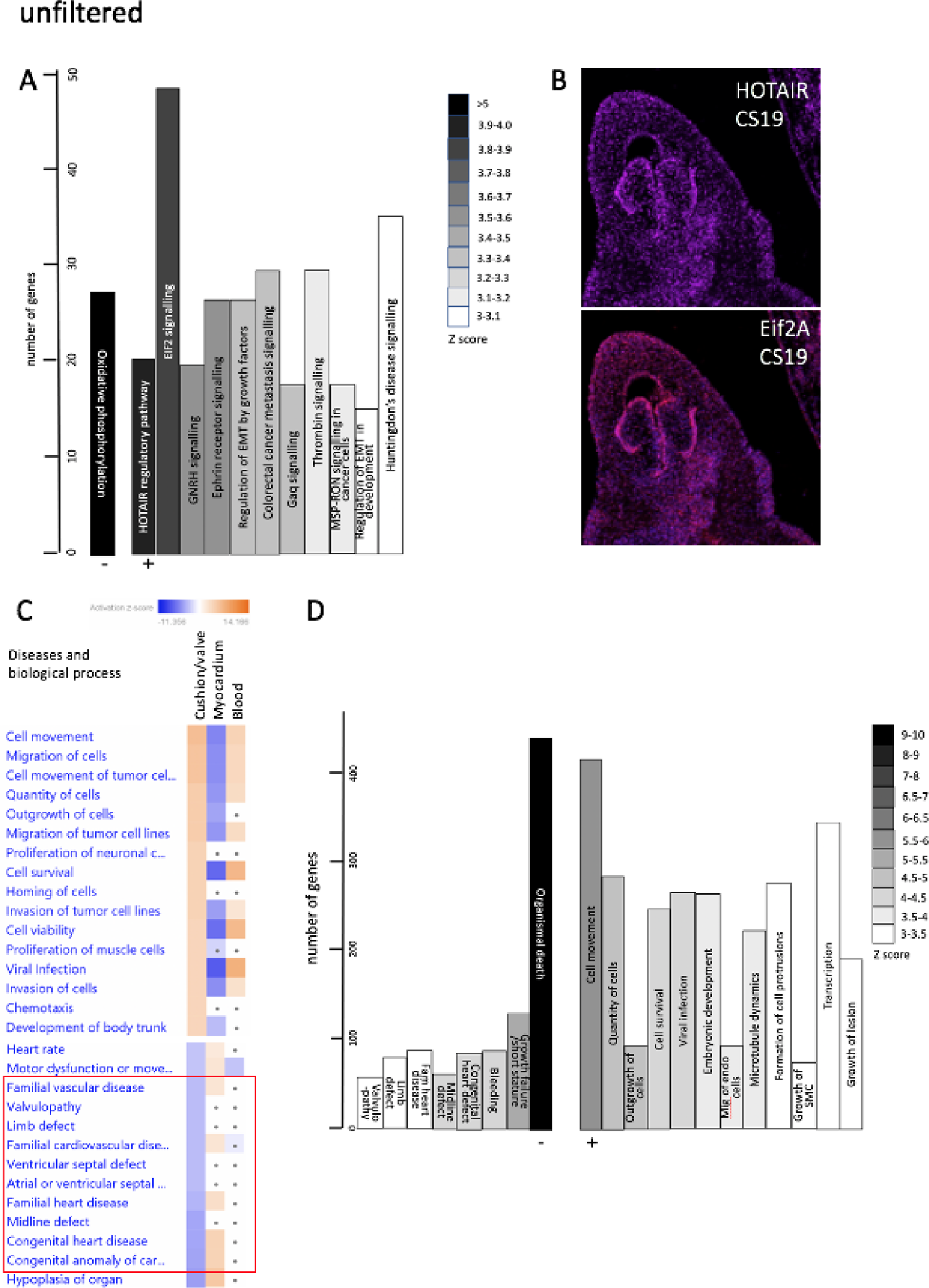
Functional associations of valve cluster genes generated by unfiltered IPA analysis. **A)** If no filtering is carried out, the HOTAIR and EIF2 signalling pathways are most strongly associated with the valve cluster data, whereas oxidative phosphorylation is negatively associated. **B)** Both HOTAIR and Eif2A are expressed in the endocardium of developing aortic valves at CS19. **C,D)** Cell movement/migration and proliferation-related terms are most strongly associated with the unfiltered valve dataset (compared with the myocardial), whereas terms relating to cell/tissue death and again congenital/familial heart/cardiovascular disease, including valvulopathy, (red box in C) were negatively associated.

### “Valve” genes are frequently associated with cardiac malformation and disease

Having identified a set of genes that were highly differentially expressed in human embryonic cardiac cushion tissue compared to the myocardium, but that were not well represented in the gene sets associated with congenital heart defects and cardiac development, we wanted to know if there was data in the literature to support an association between our gene list and heart malformation. We carried out a literature search using OMIM and PubMed to identify data linking the top 100 differentially expressed cushion genes and either human congenital heart disorders or cardiac defects in mouse knockouts (Supplementary Table 5). Breaking this data into quartiles, 15/25 genes in the top quartile were associated with congenital heart defects or diseases affecting the outflow region in mouse or human, compared with only 4/25 in the bottom quartile. We looked in more detail (using literature searches and interrogation of the Mouse Genome Informatics database; https://www.informatics.jax.org) at the phenotype of mouse knockouts of the novel genes LGALS1, PDLIM3, THBS1, LINC00632, RBP1, FLRT2, OLFML3, SLN, MXRA8, S100A11 and C7 that we highlighted earlier. Of these, there was no reported cardiac phenotype for LGALS1, OLFML3, MXRA8 and S100A11 mouse knockouts (Poirier and Robertson, 1993; Ikeya et al, 2005). PDLIM3, THBS1 and FLRT2 knockout mice are reported to have cardiac defects although the valves were not specifically described (Pashmforoush et al, 2001; Lawler et al, 1998; Muller et al, 2011). SLN knockouts are reported to have abnormalities in cardiac contractility (Babu et al, 2007) but no congenital abnormalities were reported, whereas C7 mice are reported to have neurological disturbances (https://www.informatics.jax.org/allele/MGI:5823284). RBP1 knockout mice are reported to have cardiac metabolic disturbances but the anatomy of the heart does not appear to have been analysed in embryos or in adult animals (Matt et al, 2005; Zalesak-Kravec et al, 2022). Thus, there is a high incidence of great artery and valve malformations and/or disease associations with the most differentially expressed genes in our valve cluster, although in several cases null mice have not been investigated for these abnormalities.

### RBP1 knockout mice have abnormalities of the arterial valves

We were intrigued that RBP1 and CRABP2 (cellular retinoic acid binding protein 2) were both found in our list of the top 30 most differentially expressed genes in the valve cluster (Figure 3A) and that RARβ was one of the top 30 upstream regulators (Figure 5A) of the valve cluster. Although the retinoic acid signalling pathway has not previously been specifically associated with arterial valve development, we examined the expression patterns of RBP1 (also known as CRBP1) and CRABP2 in more detail in both mouse and human embryos. RBP1 was specifically localised to the arterial valve endocardium at all stages examined (E11.5-E13.5 in mouse, CS16-21 in human) in both mouse and human embryos, whereas CRABP2 was found in the surrounding arterial wall in both species (Figure 7A).

**Figure 7.**
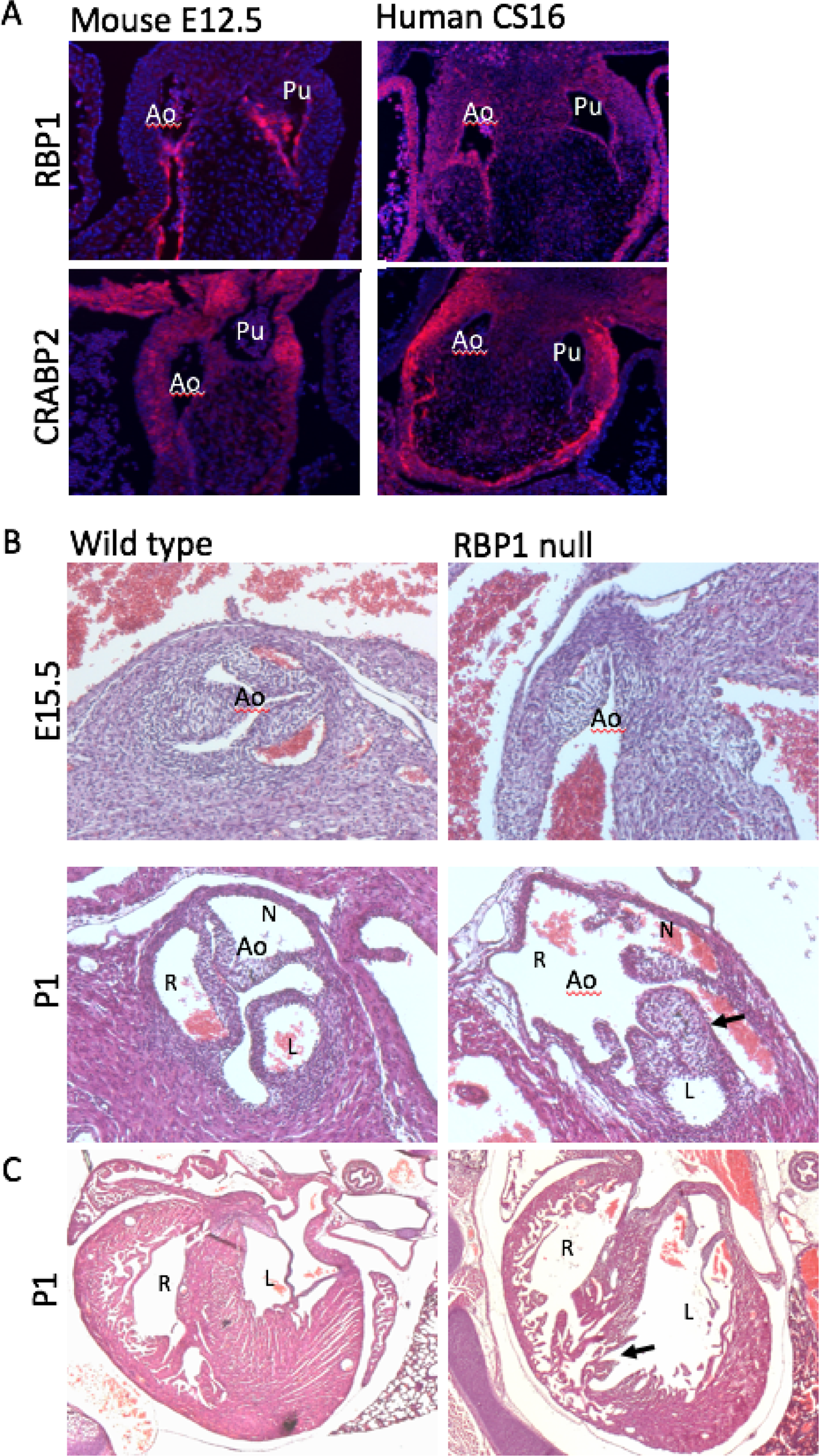
RBP1 plays a crucial role in the developing arterial valves. **A)** RBP1 is expressed (pink dots) in the endocardium of the developing aortic and pulmonary valves in mouse E12.5 and human CS16 hearts. In contrast, CRABP2 expression (pink dots) localises to the wall surrounding the valve leaflets in both species at the same timepoints. **B)** Abnormalities of the aortic valve including BAV (in the E15.5 mutant) and valve dysplasia (arrow in P1 mutant) were seen in late fetal and neonatal RBP1 mutants. **C)** RBP1 mutant neonates also have abnormalities on the ventricular myocardium including muscular ventricular septal defects (arrow). Stage-matched and oriented control aortic valves are shown for comparison. Ao = aortic valve; L = left; N = non-coronary; Pu = pulmonary valve; R = right.

Whereas the mouse model for CRABP2 has been described in detail, with only mild digit defects reported (Lampron et al, 1995), the embryonic phenotype of RBP1 knockout mice does not appear to have been analysed in any detail (Matt et al, 2005; Zalesak-Kravec et al, 2022). We therefore obtained RBP1 knockout fetuses and neonates at E15.5 and P1 respectively and analysed the hearts for structural heart defects. Four RBP1 knockout fetuses were examined at E15.5 and compared with wild type controls on the same genetic background. A range of defects affecting the ventricular myocardium and the outflow region of the heart were apparent (Figure 7B). These included abnormalities in the positioning of the great arteries so that the arteries were parallel in 2/4 mutants analysed, and a tortuous pulmonary trunk in 3/4. Moreover, BAV was found in 2/4 RBP1 knockouts examined at E15.5. Four RBP1 knockouts were also examined at P1, with a similar spectrum of defects observed (Figure 7 B,C). Arterial valve defects were seen in 3/4 mutants with BAV in one and dysplastic or abnormal valve attachments seen in 2/4. Ventricular septal defects affecting the muscular part of the septum were also a common finding, affecting 4/8 of the E15.5 and P1 mutants examined.

## Discussion

ST is becoming more popular as a technique for gathering spatially-related transcriptomic data (DeLaughter et al, 2016; de Soysa et al, 2019; Asp et al, 2019). The potential to measure transcript expression levels whilst maintaining the spatial context from which they arose, together with the new insights into cellular organisation this data reveals, is an obvious biological research goal with many large-scale consortia (e.g. Human cell Atlas, Human Developmental Biology Initiative) utilising ST as a key technology to realise this aim. However, ST, as a method, is still in its infancy. As such, no clear recommendations on best practice have yet emerged. One key aspect of the method that has yet to be resolved is the number of replicates required to produce robust datasets. Although we did not see major differences between our technical and biological replicates it is important to remember that, as with any transcriptomic study, the number of replicates influences the statistical power of the experiment (McIntyre et al, 2011). Biological replicates are needed to detect differentially expressed genes between conditions, whereas technical replicates provide a means to distinguish true differences in gene expression from experimental artefacts. We have taken our lead from that of single cell RNA sequencing and have adopted an approach of using at least three technical replicates (tissue sections) for each tissue sample. This strategy acts to increase the number of spots for each given spatial position and therefore increases the transcriptional data obtained at that position, thereby statistically powering up the assay. In an ideal world, a similar approach would be taken for the biological replicates, with a suggestion that a minimum of three samples at each specific sampling time point should be used. However, it is acknowledged that obtaining rare patient samples, or as in this study, early human embryos, is often challenging. Even with the increased resolution of new ST technologies (e.g. 10X Genomics Visium, see below), technical replicates will remain an important consideration, particularly for studies where the area of interest is small or small cell populations fall between the spot capture areas, as we saw with our analysis of the valve leaflets. It will be interesting to see how the community responds to address the question of the number of replicates required for confident analysis of the data as more researchers adopt ST.

The technologies for performing ST are continuing to be developed and improved upon and there are now multiple commercial ST applications available. Indeed, the method used here has now been superseded by release of the 10X Genomics Visium kits which have doubled the spot resolution. Regardless, the data presented here is still very much of value, especially given the limited number of other outflow cushion-specific human datasets and given the validation by *in situ* methods and overlap with existing human and mouse data. Despite the lower resolution of our study, we found that our use of technical replicates provided us with enough datapoints to distinguish three spatially distinct biological regions. However, even with these replicates we were unable to study individual valve leaflets in detail. Statistically significant data between cushion-derived and intercalated valve swelling-derived valve leaflet precursors was not achieved and points towards higher resolution studies being required to address this when the technology is capable of single cell resolution in the future. Such advancements in ST are already underway and the spatial resolution of the technology can, or very soon will, be able to resolve to a scale smaller than that of a single cell (from review see Cheng et al, 2023). However, this does not on its own mean that single cell resolution can be achieved for ST from tissue sections. It is extremely difficult to section any tissue uniformly to a single cell thickness and therefore the data sampling points are always likely to contain transcriptomes from more than one cell. The sequencing costs involved in the scale up to one cell resolution are also likely to be prohibitive. It is perhaps expected therefore, that the move to single cell resolution for ST will come from a combination of increased resolution of the ST assay in conjunction with advances in computational integration of scRNA-seq and ST data - the single cell resolution coming from the data obtained from sc-RNA seq and the spatial positioning from ST. Strides have already been made in this regard with software packages such as Cell2location, MERINGUE, and SquidPY, (Miller et al, 2021; Kleshchevnikov et al, 2022; Palla et al, 2022) already allowing for such combined analysis and new applications enabling more sophisticated analysis are anticipated.

Comparison between the transcriptome of the CS16 and CS19 hearts showed that the two datasets showed a high degree of similarity, although separated by approximately 8-10 days. A related study using human embryos (Asp et al, 2019) came to a similar conclusion and merged data from 4-5-9 post conceptional weeks. Although the core genes expressed were very similar at CS16 and CS19, there were on average more than 50% more genes captured at CS19 compared to CS16. Between these Carnegie stages the outflow tract is actively growing and undergoing septation, although these processes are not complete for another 3-5 days of gestation beyond CS19. Thus, although heart morphology changes over this time frame, the genes regulating and supporting this morphogenesis overlap to a high degree. Three non-overlapping (in terms of spots from CS16 and 19 hearts) clusters were identified at these time points. Analysis of the most differentially expressed genes, as well as “perfect marker” analysis, confirmed these as myocardial, valve and blood (in the lumen of the heart). Notably, we could not establish a distinct endocardial dataset, with genes for this cell type found in all 3 clusters. This likely reflects the large relative size (100μm) of the spots, which is larger than the typical thickness (10-30μm) of the endocardial monolayer. More recent versions of the technique have significantly better resolution (55μm) although it is likely that resolution of the endocardial monolayer will continue to be a problem until single cell resolution is achieved. Strikingly, 29/30 of the top differentially expressed genes in the myocardial cluster, and 28/30 of the genes in the blood cluster, were already known to be abundantly expressed in these tissues. In contrast, 11/30 of the differentially expressed genes in the valve cluster had not previously been linked to the cardiac valves, at least until transcriptomic studies came along (see Figure 3), suggesting that this tissue is less well annotated at the transcriptional level than are myocardium or blood. This would appear to be supported by the fact that IPA analysis showed that the genes in the valve cluster were negatively associated with terms linked to congenital heart disease and familial heart disease anomalies (Figure 5). In contrast, our myocardial cluster was strongly correlated with these terms. This surprising result suggests that datasets linked to congenital heart disease/familial heart disease over-represent genes expressed in cardiomyocytes (which are the major cell type in the developing heart), compared to those expressed in valve tissue. For large genomic studies, some degree of filtering is essential in order to whittle down the huge number of variants detected to a workable number for further analysis. The unintended outcome from this is that mostly cardiomyocyte genes will be positively selected when BAV/CHD genomic data are filtered for “cardiac development” terms, because valve-related genes are under-represented in the relevant datasets. As a consequence, potential causative variants may be excluded from further analysis. Without wanting to over interpret these results, this may explain why relatively few causative genes have been identified by genomic studies of CHD cases, and even fewer have been validated as being disease causing.

Extracellular matrix molecules, cytoskeletal proteins and transcriptional regulators were the commonest classes of protein encoded by the most abundant genes in the valve cluster, participating in cellular processes such as EMT (EndMT in the cushions/valves), cell movement/migration and proliferation – all processes that are known to be crucial in the remodelling valves (reviewed in Henderson et al, 2020, deVlaming et al, 2013) and other reviews). Importantly, our data overlaps and complements the existing datasets for developing valves generated from mouse tissue from a range of embryonic and postnatal stages. For example, genes associated with bone formation were highly represented in our valve dataset, as has been reported for mid-late fetal mouse valves (Chakraborty et al, 2008). Similarly, considerable overlap was found with our dataset and single cell RNASeq data from postnatal mouse valve interstitial cells (Hulin et al, 2019), Comparisons with the first trimester human outflow tract and atrioventricular valves (Asp et al; 2019), with 25-30% overlap between the two datasets for the top 250 genes in each. Thus, our data is well supported by data generated from other studies and adds an important transcriptional dataset representing the developing human arterial valves.

Variants in several of the most differentially expressed genes in the valve cluster, including COL1A2, ELN and FBLN1 (Callewaert et al, 2011; Wu et al, 2021; Theis et al, 2022), have been suggested as being potentially causal for BAV in human genomic studies, whereas MFAP4 and GPC3 have been implicated as causal genes for other CHD (Hitz, 2012; Tomita-Mitchell et al, 2012). Notably though, COL1A2, ELN and FBLN1 are all expressed in the arterial wall surrounding the valve (Figure 3) and are primarily implicated as causal genes for aortopathy, with their roles as causal genes for BAV unproven. Although several of the highly differentially expressed genes we identify have not previously been shown to be expressed in the developing valve region, almost all, where expression data was available, were confirmed as being expressed in the arterial roots where the valves are located. Thus, the ST appears to have successfully identified the arterial valve transcriptome at the CS16-19 stage of human development. Four genes (STX10, HES4, MXRA5 and LINC00632) in the top 250 differentially expressed genes in the arterial valve cluster are not found in the mouse genome and therefore could not have been detected from mouse-based transcriptomic studies. The expression patterns of these genes all confirmed expression in the arterial roots, with MRXA5 and LINC00632 found in the forming valve leaflets. HES4 is an important downstream target of Notch signalling and has been identified as being differentially expressed in VIC from calcific aortic valve disease (Xu et al, 2020). MRXA5 has been reported to be expressed in cushion tissue in the chicken embryo (Robbins and Capehart, 2018) although nothing (until now) was known about its expression pattern in developing human heart. All of these genes can now be considered potential candidates for human arterial and valve disease.

IPA analysis, whether filtering the data for cardiac and development, or without this filter, highlighted CXCL12, TGFBR1 and more generally, pathways involved in regulating EMT, in the remodelling phases of valve development. As already mentioned, EndMT is a key process in cushion/valve development and this process is initiated by the TGFβ signalling pathway (reviewed in deVlaming et al, 2013). Although EndMT should be mostly completed by CS16 (equivalent of mouse E12.5), most of these genes continue to play roles during the remodelling stages of valve development (reviewed in deVlaming et al, 2013; Henderson et a, 2020). CXCL12 has also been shown to play an early role in the directional migration that is a crucial part of EndMT, but also in regulating cell proliferation at later remodelling stages (Ridge et al, 2021). Similarly, TGFBR1 (ALK5) also plays crucial roles in valve development (Mercado-Pimental et al, 2011). The unfiltered analysis also suggested that HOTAIR and EIF2 signalling may play important roles in valve development, although there is nothing known about these pathways in the developing heart. We were particularly intrigued by the HOTAIR pathway as this is a long non-coding RNA that regulates HOX gene expression (Rinn et al, 2007). HOXA3 was highly differentially expressed in the valve cluster, appearing in the top 30 transcriptional regulators (Supplementary Figure 6B). Interestingly, null mutants for HOXA3 (then called hox1.5) are reported to have cardiac defects including BAV (Chisaka and Cappechi, 1991), confirming that this gene is vital for arterial valve development. HOX genes are generally thought to be regulated by retinoic acid signalling in the developing heart (Bertrand et al, 2011; Zaffran and Kelly, 2012), so we were interested that two genes in this pathway, RBP1 and CRABP2, were amongst the top 30 differentially expressed genes in the valve cluster, with immunohistochemistry confirming their localisation to the developing arterial valve leaflets in the human and mouse. Moreover, RARβ was predicted to be one of the most important regulators of genes in the cluster. Although there is considerable evidence that retinoic acid plays an important role in endocardial cushion development, its precise roles remain unclear. Excess and deficient RA signalling disrupts formation of the endocardial cushions (Yan and Sinning, 2001; Gruber et al, 1996), at least in part through the dysregulation of key genes, such as Tbx2 (Sakabe et al, 2012). RA has also been shown to be critical for the development of the coronary vasculature (Wang et al, 2018). Given these findings, it seems that RA levels need to be tightly controlled for normal development to proceed. Analysis of RBP1 knockout fetuses and neonates revealed a range of defects affecting the outflow of the heart, particularly BAV and dysplastic arterial valve leaflets. Thus, RBP1, and presumably RA signalling more generally, appears to be crucial for development of the arterial valve leaflets and thus are potential causative genes for BAV and other valve malformations.

This study has shown the utility of ST for generating arterial valve-specific transcriptomic data from human embryos, broadening our knowledge of the genes and signalling pathways important in human valve development, and acting as a foundation for more targeted screening of variants for common arterial valve malformations such as BAV.

## Materials and Methods

### Human and mouse embryos

Human embryos were obtained from the Human Developmental Biology Resource (HDBR) following social terminations of pregnancy with informed donor consent and ethical approval from the Newcastle and North Tyneside 1 NHS Health Authority Joint Ethics Committee (08/H0906/21+5). Heart samples, Carnegie stage (CS)16 and CS19, were dissected from the embryo, embedded into Optimal Cutting Temperature (OCT) and snap-frozen in an isopentane bath on liquid nitrogen. Sections were taken using a Leica cryostat at 10μm intervals, RNA extracted from four consecutive tissue sections and RNA Integrity Number (RIN) analysis performed using an Agilent Bioanalyzer. Samples with a RIN value of 7 or above were deemed suitable for ST analysis.

Mouse embryos (CD1 or C57Bl/6) were obtained from timed matings carried out overnight, with the presence of a copulation plug designated embryonic day (E) 0.5. *Rbp1*^-/-^ mice (C57BL/6N background) were bred similarly according to institutional guidelines of the University of Maryland, Baltimore. *Rbp1*^-/-^ mice were originally obtained from Pierre Chambon and Norbert Ghyselinck (Institut de Genetique et de Biologie Moleculaire et Cellulaire, Institut National de la Santé et de la Recherche Medicale, Illkirch, France). Mice were maintained according to the Animals (Scientific Procedures) Act 1986, United Kingdom, under project license P9E095FF4. All experiments were approved by the Newcastle University Ethical Review Panel.

### Spatial transcriptomics

An overview of the ST methodology and workflow employed is shown in Figure 1C. 10μm tissue sections were collected into the capture windows of an ST Tissue Optimisation slide (10X Genomics cat. no. 1000131) and the permeabilisation conditions for the ST analysis of the heart tissue determined following the method described in the Tissue Optimisation Manual (Spatial Transcriptomics, version 190219). Briefly, this involved pre-permeabilisation of the sections in 50U/μl collagenase at 37°C for 20 minutes, followed by permeabilisation using a staggered sequence of 0-12 minute intervals in 0.1% pepsin/HCl at 37°C. RNA captured by the oligo-dT probes bound to the slide was reverse transcribed and Cy3-labelled nucleotides incorporated into the reaction. The optimal permeabilisation time was then determined by visual inspection of the resultant fluorescent cDNA footprint on each section.

Consecutive tissue sections in the regions containing the aortic valves (5 at each time point) were then sectioned to an ST Library Preparation slide (10X Genomics, cat # 1000132), H&E stained (Supplementary Figure 1), and the tissue sections and underlying spots (each spot covering a 100μm region and separated by distance of 200μm from the centre of each neighbouring spot) captured using a Nikon TiE inverted microscope at 10x magnification (Plan Fluor 10x 0.3 NA). Raw images were stitched together using NIS-Elements software with parameters set to 20% overlap and stitching via optimal path method. ST analysis was performed as per the manufacturer’s instructions (Spatial Transcriptomics Library Preparation Manual, version 190219). Following enzymatic cleavage of the barcoded probes and associated captured RNA transcripts from the slide, sequencing libraries were generated and sequenced at 50,000 reads per sampling region (spot) using an Illumina NovaSeq 6000.

### Alignment and Quantification

The paired FASTQ files were de-multiplexed with a publicly available perl script (https://github.com/tallulandrews/scRNASeqPipeline/blob/master/0_custom_undo_demultiplexing.pl) using the spatial barcodes encoded in read 1. Read 2 from successfully de-multiplexed pairs were trimmed for quality using Trimmomatic version 0.3 (Bolger et al, 2014). A reference was created using Ensembl human reference genome (GRCh38.p12). The trimmed reads were aligned to the reference using STAR version 2.5.3a in single read alignment mode (Dobin et al, 2013). The number of reads were quantified using HTSEQ version 0.6.1 (Anders et al, 2015) and a count matrix was created. Spot detector software (Wong et al, 2018) was used to align the fluorescent image with the brightfield image to determine the pixel co-ordinates for the spots.

### Quality Control and Visualisation using Spaniel R package

We developed the Spaniel R package (Queen et al, 2019) which provides a framework for quality control, visualisation and pre-processing of ST data. Spaniel is available from bioconductor (https://www.bioconductor.org/packages/release/bioc/html/Spaniel.html) and is built on existing S4 objects namely the SingleCellExperiment object and Seurat object to facilitate the use of other single cell methodologies. The gene expression matrix was imported in R using the Spaniel package and quality control steps were performed to remove spots which fell outside the tissue area. We assessed the number of genes per spot, number of reads per spot and percentage of mitochondrial reads per spot. High levels of mitochondrial genes can indicate poor sample quality leading to abnormal gene expression patterns caused by the capture process rather than underlying biology of the sample itself. Samples with low quality are also likely to have low numbers of gene or reads detected. Section 2 of the CS16 tissue was removed from analysis because of quality control issues, leaving 4 sections for analysis at CS16 and 5 at CS19.

### Clustering analysis and Visualisation of Gene Expression

Gene expression was normalised and highly variable genes were identified using Seurat. We firstly combined the technical replicates from the adjacent tissue sections into one dataset using Seurat integration methods to account for batch effects from the data (Stuart et al, 2019). Secondly, the spots from all sections at each time point were combined into one dataset, again using Seurat integration methods. Clustering analysis was performed on each of the integrated datasets with a resolution of 0.8. Differential Expression analysis was performed to identify changes in gene expression between the clusters using the Wilcoxon test.

### IPA analysis

Pathway analysis was performed using Qiagen Ingenuity Pathway Analysis (IPA) software. The marker gene lists for each of the three clusters, with an adjusted p value of less than 0.05 were run through the core analysis module of IPA. A comparison analysis was then performed to look for differences and similarities between the canonical pathways, upstream regulators and disease terms associated with each set of marker genes. A significance threshold of an absolute value of 2 for the z score was set where a z-score of 2 or higher indicates that a pathway is activated and –2 or lower indicates that is inhibited. The pathways were filtered for just those containing the terms cardiac or development (using the wildcard *card* or *dev *). Network plots were generated using IPA software.

### Fluorescent Immunohistochemistry

Methodology was as published previously (Eley et al, 2018). Sections were cut from paraffin-embedded embryos at 8 µm using a rotary microtome (Leica). Slides were de-waxed with Histoclear (National Diagnostics) and rehydrated through a series of ethanol washes. Following washes in PBS, antigen retrieval was performed by boiling slides in citrate buffer (0.01 mol/L) pH 6.3 for 5 minutes. Sections were blocked in 10% FCS and then incubated overnight at 4°C with the following primary antibodies diluted in 2% FCS: LGALS1 (Abcam ab138513), RBP1 (Invitrogen PA528713), CRABP2 (Abcam ab211927). After washing, sections were incubated at room temperature for one hour, with secondary antibodies conjugated to Alexa 568 (Life Technologies). Fluorescent slides were then mounted with Vectashield Mounting medium with DAPI (Vector Labs). Immunofluorescence images were collected with using a Zeiss Axioimager Z1 fluorescence microscope equipped with Zeiss Apotome 2 (Zeiss, Germany). Acquired images were processed with AxioVision Rel 4.9 software.

### RNAScope

All reagents used were taken from the RNAscope Multiplex Fluorescent Reagent Kit v2 (bio-techne, cat# 323100). 8µm sections were cut from formalin-fixed paraffin-embedded embryos using a Leica microtome. The slides were heated to 60°C for 10 min and allowed to cool to room temperature before dewaxing in xylene, rinsing in two changes of 100% ethanol and air drying. Following incubation in hydrogen peroxide at room temperature for 10 minutes and rinsing successive changes of water, the slides were then submerged in Target Retrieval solution and placed in a boiling water bath for 20 minutes. After rinsing the slides in 100% ethanol and air drying, protease plus solution was added to the sections for 30 minutes at 40°C. RNAscope probes were diluted into either probe diluent (bio-techne, cat # 300041) or a channel 1 probe as appropriate in a 1 in 50 volume, and the slides incubated at 40°C for 2 hours. Following sequential amplification steps with AMP1 and AMP2 solutions, the slides were incubated with HRP-C1 at 40°C for 15 minutes, rinsed in wash buffer and incubated at 40°C for 30 minutes in OPAL flurophore 520 (Akoya Biosciences) diluted 1 in 100 with TSA diluent. After blocking for 15 minutes at 40°C with HRP blocker, the same steps were repeated with HRP-C2 solution and OPAL flurophore 570 (Akoya Biosciences). After nuclei staining with DAPI, slides were mounted in Prolong Gold Antifade (Thermo Fisher Scientific) and imaged as above. Human RNAscope probes used in this study (bio-techne): PDLIM3 (cat. # 533411-C3), ID4 (cat. # 466371), STX10 (cat. # 1119651), HES4 (cat. # 548621-C3), MRXA5 (cat. #419691-C2), HOTAIR (cat. # 312341-C2) and EIF2A (cat. # 410701).

### BaseScope

8 µm sections were cut from formalin-fixed-embedded embryos using a Leica microtome. Slides were heated to 60°C and allowed to cool to room temperature. Slides were then dewaxed in two changes of xylene (5 minutes each), rinsed in two changes of 100% ethanol followed by air drying. Sections were covered with hydrogen peroxide and incubated at room temperature for 10 minutes and rinsed in two changes of diethyl pyrocarbonate (DEPC) treated water. Following incubation and rinsing, slides were then placed in boiling target retrieval solution for 20 minutes, then immediately rinsed in two changes of DEPC water. After rinsing, slides were rinsed in two changes of 100% ethanol and allowed to air dry. Protease IV was added to cover sections, and incubated for 30 minutes at 40°C. Following rinsing in 2 changes of DEPC water, two drops of channel 1 BaseScope probe BA-Hs-LINC00632-213-2zz-st-C1 (bio-techne, cat# 1128991-C1) was added to sections and incubated at 40°C for 2 hours. Following 8 sequential amplification steps with AMP1 – AMP8, 1:60 BaseScope Fast RED-B was added to BaseScope Fast RED-A and pipetted onto sections (BaseScope v2 Red Assay - bio-techne, cat# 323910). After a 10 minute incubation period, slides were rinsed in DEPC water and counterstained with 50% methyl green solution (Vector, Cat# H-3402). Slides were then heated to 60°C for 30 minutes, mounted with EcoMount (Vector, cat# H-5000), coverslip and imaged as above.

## Acknowledgements

The human embryonic and fetal material was provided by the Joint MRC/Wellcome (MR/R006237/1, MR/X008304/1 and 226202/Z/22/Z) Human Developmental Biology Resource (www.hdbr.org).

